# PD-1 negatively regulates helper T cell differentiation into Th2

**DOI:** 10.1101/2024.07.26.605269

**Authors:** Masaki Tajima, Naoko Ikuta, Yuka Nakajima, Kensuke Suzuki, Yosuke Tokumaru, Peng Li, Hiroshi Kiyonari, Tasuku Honjo, Akio Ohta

**Affiliations:** Department of Immunology, Institute of Biomedical Research and Innovation, Foundation for Biomedical Research and Innovation at Kobe, Kobe 650-0047, Japan; Division of Integrated High-Order Regulatory Systems, Center for Cancer Immunotherapy and Immunobiology, Graduate School of Medicine, Kyoto University, Kyoto 606-8507, Japan; Pharmaceutical R&D Division, Meiji Seika Pharma Co. Ltd., Tokyo 104-8002, Japan; Laboratory for Animal Resources and Genetic Engineering, RIKEN Center for Biosystems Dynamics Research, Kobe 650-0047, Japan; Department of Immunology and Genomic Medicine, Center for Cancer Immunotherapy and Immunobiology, Graduate School of Medicine, Kyoto University, Kyoto 606-8507, Japan

## Abstract

Programmed Cell Death Protein-1 (PD-1) represents endogenous mechanisms of negative immunoregulation. While the modulation of effector functions has been the major focus of PD-1 research, quick PD-1 upregulation in naïve T cells starting 1 h after priming raised a possibility that PD-1 also affects the development of effector T cells. The role of PD-1 in functional differentiation into Th1 and Th2 has been unclear. In murine naïve CD4^+^ T cell activation, we found that PD-1 stimulation during the early stage of T cell activation strongly impaired Th2 cell development, while Th1 cell induction was relatively resistant to this immunosuppressive signaling. The steep decline in Th2 cell induction suggested the significance of PD-1 in allergic inflammation. Treatment with anti-human PD-1 agonist antibody inhibited allergic inflammation in human PD-1-knock-in mice as shown by the reduction of Th2 cells, IgE levels and eosinophilic infiltration. This study shows that PD-1 regulates not only the intensity but also the quality of immune response by deviating Th differentiation. PD-1 stimulators are projected to be valuable in suppressing various forms of inflammatory activities, but the efficacy against Th2-dominant immune response may be particularly high.

## Introduction

Th2 cells, which are CD4^+^ T cells secreting IL-4, IL-5 and IL-13, drive allergic inflammation as a major component of type 2 immune response (*1, 2*). Overactive type 2 immune response is responsible for the typical allergic diseases involving IgE-mediated degranulation of mast cells. IL-4 promotes B cell response and antibody class switch to IgE. IL-13 was shown to cooperate with IL-4 to produce high-affinity IgE, which is superior in stimulating mast cells than low-affinity IgE (*3*). IL-5 and IL-13 can promote allergic inflammation in an IgE-independent manner by activating eosinophils. These cytokines may be also produced from other effectors of allergic inflammation such as innate lymphoid cells (ILC) 2, mast cells and basophils, and mediate a positive feedback loop expanding type 2 immunity (*4, 5*). The growing prevalence of allergic diseases has become a worldwide major health concern. The prevention of overactive type 2 immune response is crucial to the control and treatment of various allergic disorders.

Recent advances in the understandings of type 2 immunity revealed that early production of IL-25, IL-33 and thymic stromal lymphopoietin (TSLP) from stressed epithelium can initiate type 2 immune responses (*2, 6*). ILC2 respond to these epithelial cell-derived cytokines and translate the danger signal into setting up the cytokine milieu that can specifically promote functional differentiation of naïve CD4^+^ T cells to Th2. However, other factors such as T cell receptor (TCR) signal intensity or co-stimulation can also affect Th2 cell differentiation (*1, 7*), and their relative importance remains to be elucidated (*8, 9*).

PD-1 represents negative co-stimulatory receptors and is indispensable in controlling T cell activities. The engagement of PD-1 with its natural ligands, PD-L1 or PD-L2, recruits Src homology-2 domain-containing protein tyrosine phosphatase-2 (SHP2), which in turn dephosphorylates the components of TCR signaling cascade and attenuates T cell activation (*10–12*). The significance of PD-1-dependent regulation has been evident from the exaggerated inflammatory activities in mice lacking PD-1 or PD-L1 (*12*). This discovery led to the emergence of PD-1 as a target of immune manipulation, and blockers of PD-1 pathway have seen a clinical success in cancer immunotherapy (*13–15*). Of note, eosinophilia and adverse allergic inflammation have been reported in some cancer patients after the treatment with PD-1 blocking antibodies (*16–19*). These symptoms imply the relevance of PD-1 pathway in modulating Th2 cells and type 2 immune response. The association of PD-1 and allergic inflammation has been studied in animal models using blocking antibodies or knockout mice of PD-1 or PD-L1; however, the results are surprisingly inconsistent. In allergic conjunctivitis and dermatitis models, the blockade of PD-1 or PD-L1 increased Th2 cells but failed to enhance inflammation (*20, 21*). For the allergic asthma induction, the blockade of PD-1 or PD-L1 did not exaggerate airway hypersensitivity or eosinophils infiltration (*22, 23*), and asthma induction in PD-L1-knockout mice rather reduced lung inflammation (*24*).

In this study, our purpose is to elucidate the role of PD-1 in Th2 cell development. PD-1 expression is not found on naïve T cells but is upregulated in activated T cells. Therefore, most PD-1 studies have focused on already established effector T cells. However, our finding of PD-1 upregulation in the early phase of T cell activation prompted us to examine the impact of PD-1 stimulation on the direction of effector T cell development. Recently, we established PD-1 agonist antibodies, which can trigger the immunosuppressive activity of PD-1 and suppress inflammatory diseases in vivo (*25*). PD-1 stimulation during the functional differentiation of naïve CD4^+^ T cells was found to strongly impair the induction of IL-4-producing Th2 cells, suggesting the feasibility of PD-1 as a target of therapeutic intervention to Th2-related diseases.

## Results

### Early PD-1 signaling substantially affects Th1/Th2 balance

PD-1 expression is not detectable in naïve T cells, but its upregulation after activation serves as a negative feedback mechanism to limit effector T cell activities. PD-1-dependent immunoregulation has been shown to critically affect anti-pathogen response and autoimmunity by activated T cells (*12*). Besides the control of effector T cell activities, we hypothesized that PD-1 stimulation may change the course of effector T cell development when taking place in the early phase of T cell activation. To examine PD-1 induction during T cell activation, naïve CD4^+^ T cells from MHC class II-restricted TCR-transgenic DO11.10 mice were stimulated with OVA_323-339_ peptide. T cells rapidly downregulated CD62L and started to upregulate PD-1 within 1 h post stimulation (Figure 1A, B). PD-1 levels progressively increased thereafter. The extent of PD-1 induction was associated with the antigen dose, and PD-L1 expression on antigen-presenting cells (APC) did not affect PD-1 induction (Figure 1C). PD-L1 on APC was suppressive to the freshly activated DO11.10 cells since the blockade of PD-L1 enhanced cumulative cell proliferation and cytokine production during first three days (Figure 1D, E). It should be noted that the optimal PD-1 upregulation was evident before the stimulated T cells started proliferation (Figure 1C, D). T cells upregulated PD-1 as soon as receiving TCR stimulation, raising a possibility that PD-1 signaling in the early stage of T cell activation can affect the direction of effector T cell development.

**Figure 1.**
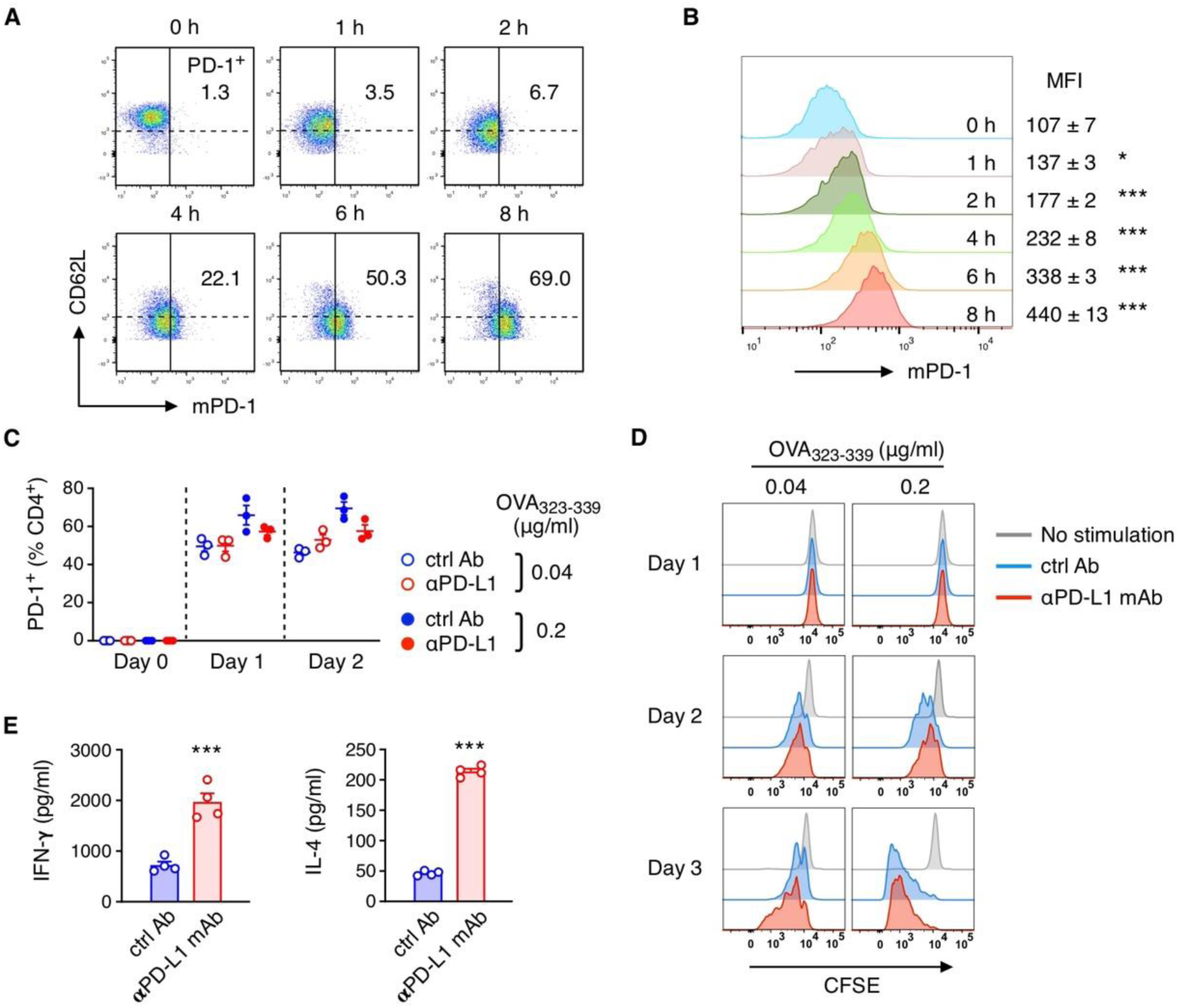
Early PD-1 upregulation in activating T cells. (A, B) PD-1 upregulation after antigenic stimulation of CD4^+^ T cells. Purified CD4^+^ CD62L^+^ T cells from DO11.10 TCR-transgenic mouse were stimulated with OVA_323-339_ peptide (1 μg/ml) in the presence of A20 cells for indicated duration of time. Numbers represent percentages of PD-1^+^ within CD4^+^ cells (A) and mean fluorescence intensity of PD-1 (B). Data represent average ± SEM of triplicate samples. (C) PD-1 upregulation on DO11.10 T cells by different concentrations of OVA peptide. Anti-PD-L1 mAb or isotype control (5 μg/ml) was added at the beginning of co-culture. (D) T cell proliferation after stimulation. CFSE-labeled DO11.10 CD4^+^ CD62L^+^ T cells were stimulated as in (C). (E) PD-1-dependent inhibition of cytokine production from DO11.10 T cells. Cytokine levels in the culture supernatant were determined after 3 days of stimulation with OVA_323-339_ (0.2 μg/ml). Data represent average ± SEM of triplicate (B, C) or quadruplicate (E) samples. *p < 0.05, ***p < 0.001; Student’s t-test.

Upon stimulation, naïve CD4^+^ T cells differentiate into functionally distinct helper subsets: Th1, Th2, Th17 or regulatory T cells among others (*26*). The intensity of T cell stimulatory signaling, which is affected by TCR affinity to the MHC complex together with numbers and duration of TCR-MHC interaction, can considerably affect the direction of naïve CD4^+^ T cell differentiation (*26, 27*). Since PD-1 stimulation disrupts TCR signaling cascade, the early PD-1 induction during T cell activation may influence functional differentiation of CD4^+^ T cells. T-bet and GATA3 are master transcription factors of differentiation and functions of Th1 and Th2 cells, respectively. Naïve DO11.10 CD4^+^ T cells undergoing functional differentiation into Th1 or Th2 upregulate either transcription factor, but the expression of T-bet and GATA3 in CD4^+^ T cells is little on day 1 compared to the expression by vast majority on day 4 (Figure 2A). In contrast, PD-1 induction was evident within hours, well before the determination for the direction of CD4^+^ T cell differentiation into Th1 or Th2.

**Figure 2.**
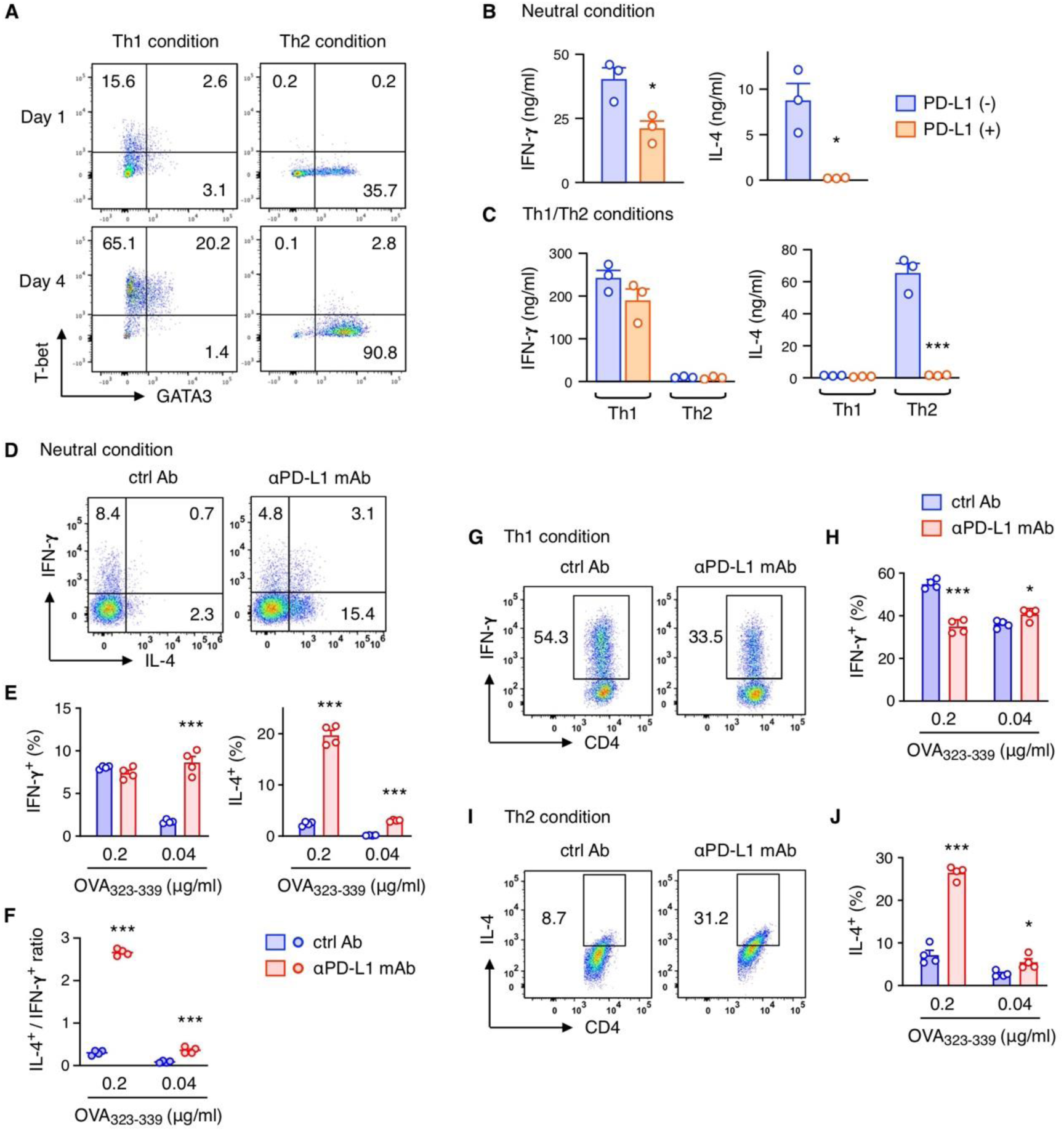
PD-L1 blockade unleashes the development of IL-4-producing Th2 cells. (A) Development of effector cells expressing T-bet or GATA3. Purified CD4^+^ CD62L^+^ DO11.10 T cells were stimulated with OVA_323-339_ (1 μg/ml) in a Th1 or Th2-skewing condition. Intracellular staining for T-bet and GATA3 expression was performed after 1 and 4 days. (B, C) PD-1-dependent regulation of Th1 and Th2 cell differentiation. Naïve DO11.10 T cells were stimulated in the neutral condition (B) or Th1/Th2-skewing conditions (C) for 3 days using PD-L1-deficient or PD-L1-expressing parental A20 cells as APC. Subsequently, cytokine-producing activities were evaluated by stimulating activated T cells (1.5 x 10^5^ cells) with immobilized anti-CD3 mAb for 24 h. (D-J) Naïve DO11.10 T cells were stimulated for 3 days using parental A20 cells as APC in the presence or absence of anti-PD-L1 mAb. T cell stimulation was conducted in the neutral (D-F) or skewing condition for Th1 (G, H) and Th2 cells (I, J). Activated T cells were restimulated in the same condition for 2 more days. Intracellular staining for cytokines was performed on day 5. Percentages of IFN-γ- and IL-4- positive events in CD4^+^ cells (D, E, G-J) and IL-4^+^ event / IFN-γ^+^ event ratios in individual samples (F) are shown. Data represent average ± SEM of triplicate (B, C) or quadruplicate (E, F, H, J) samples. *p < 0.05, ***p < 0.001; Student’s t-test.

To examine the direct impact of PD-1 stimulation during the early phase of functional differentiation, DO11.10 naïve CD4^+^ T cells were primed by PD-L1-expressing or PD-L1- deficient APC. After 3 days, the same number of activated DO11.10 cells were restimulated with immobilized anti-CD3 mAb for the evaluation of cytokine-producing activity. CD4^+^ T cell activation in the presence of PD-1 stimulation induced effector T cells that produce impaired levels of cytokines. Of note, IL-4-producing cells were totally disappeared, while 50% of IFN-γ production was still present (Figure 2B). This result is not due to the inhibition of effector function but represents the impaired establishment of cytokine-competent effector T cells because PD-L1 was present only at the priming phase. The greater degree of inhibition for IL-4-producing cells than IFN-γ-producers suggests that PD-1 stimulation during the priming phase critically affects Th2 cell development. When DO11.10 cells were skewed into Th1 or Th2 in the specific cytokine environments, the priming with PD-L1- expressing APC strongly impaired the development of IL-4-producing Th2 cells, but not IFN-γ-producing Th1 cells (Figure 2C).

The experiment above used two different cell lines, PD-L1-exprerssing and PD-L1-deficient B lymphoma, as APC. To exclude the possibility that different intensity of T cell stimulation by these two lines might have influenced the Th1 vs Th2 differentiation, we further compared CD4^+^ differentiation by PD-L1-expressing APC in the presence or absence of PD-L1 blocking antibody. DO11.10 T cell stimulation in the “neutral” condition where no cytokine milieu was manipulated except for IL-2 supplementation resulted in preferential Th1 induction. However, IL-4-producing Th2 cells prominently arose by PD-L1 blockade (Figure 2D, E). Consequently, Th1/Th2 ratio significantly shifted toward Th2 by PD-L1 blockade, suggesting that PD-1 stimulation is strongly suppressive to Th2 cell development over Th1 (Figure 2F). The same trend was observed in the enforced Th1- and Th2-skewing conditions. The significant increase of IL-4-producing cells by PD-L1 blockade indicated that early PD-1 signaling was strongly suppressive to Th2 cell induction, while Th1 development remained intact (Figure 2G-J). PD-1-PD-L1 interaction was suppressive to the proliferation of both Th1 and Th2 cells, suggesting that early PD-1 stimulation made a qualitative difference in the functional differentiation of CD4^+^ T cells.

### PD-1 agonist antibody suppresses cytokine-producing Th2 cell induction

PD-1-PD-L1 interaction during functional differentiation of T cells was inhibitory to the subsequent development of Th2-type effectors. We next examined whether pharmacological PD-1 stimulatory agent could negatively modulate Th2 responses. The clinical success of immunoenhancing anti-PD-1 blocking antibodies in cancer treatment has indicated that PD-1 is a proven target of pharmacological modulation of the human immune system (*15*). To establish a new class of immunosuppressant, we clarified the specific requirements to anti-PD-1 agonist antibodies that can trigger the immunosuppressive activity of PD-1 (*25*). One of the most effective anti-human PD-1 (hPD-1) agonist antibodies, HM266, was used to stimulate PD-1 in the course of CD4^+^ T cell differentiation.

To utilize anti-hPD-1 agonist antibodies in the antigen-specific activation of mouse CD4^+^ T cells, hPD-1 gene was retrovirally transduced into DO11.10 CD4^+^ T cells. Besides normal hPD-1 (hPD-1 WT), we also used mutant hPD-1 that is unable to transduce intracellular signaling due to the tyrosine to phenylalanine mutations in the ITIM and ITSM motifs in the cytoplasmic domain (hPD-1 YFYF) (*10, 11*). DO11.10 cells expressing either hPD-1 WT or hPD-1 YFYF at equivalent levels were skewed into Th1 or Th2 cells in the specific cytokine environment (Figure 3A). HM266 strongly inhibited the establishment of IL-4-producers from hPD-1 WT-expressing cells in Th2-inducing condition, whereas the development of IFN-γ-producing Th1 cells remained intact (Figure 3B, C). The inhibitory effect of PD-1 agonists on Th2 development was more potent when CD4^+^ T cells were stimulated at a low antigen dose than a high dose. The inverse correlation between antigen dose and Th2 inhibition suggests that the balance between the stimulatory TCR signaling and the inhibitory PD-1 signaling during early T cell activation is critical to the function of resulted effector cells. HM266 did not affect the differentiation of hPD-1 YFYF-expressing T cells, confirming that the inhibitory effect of HM266 was mediated by PD-1 signaling (Figure 3D-G).

**Figure 3.**
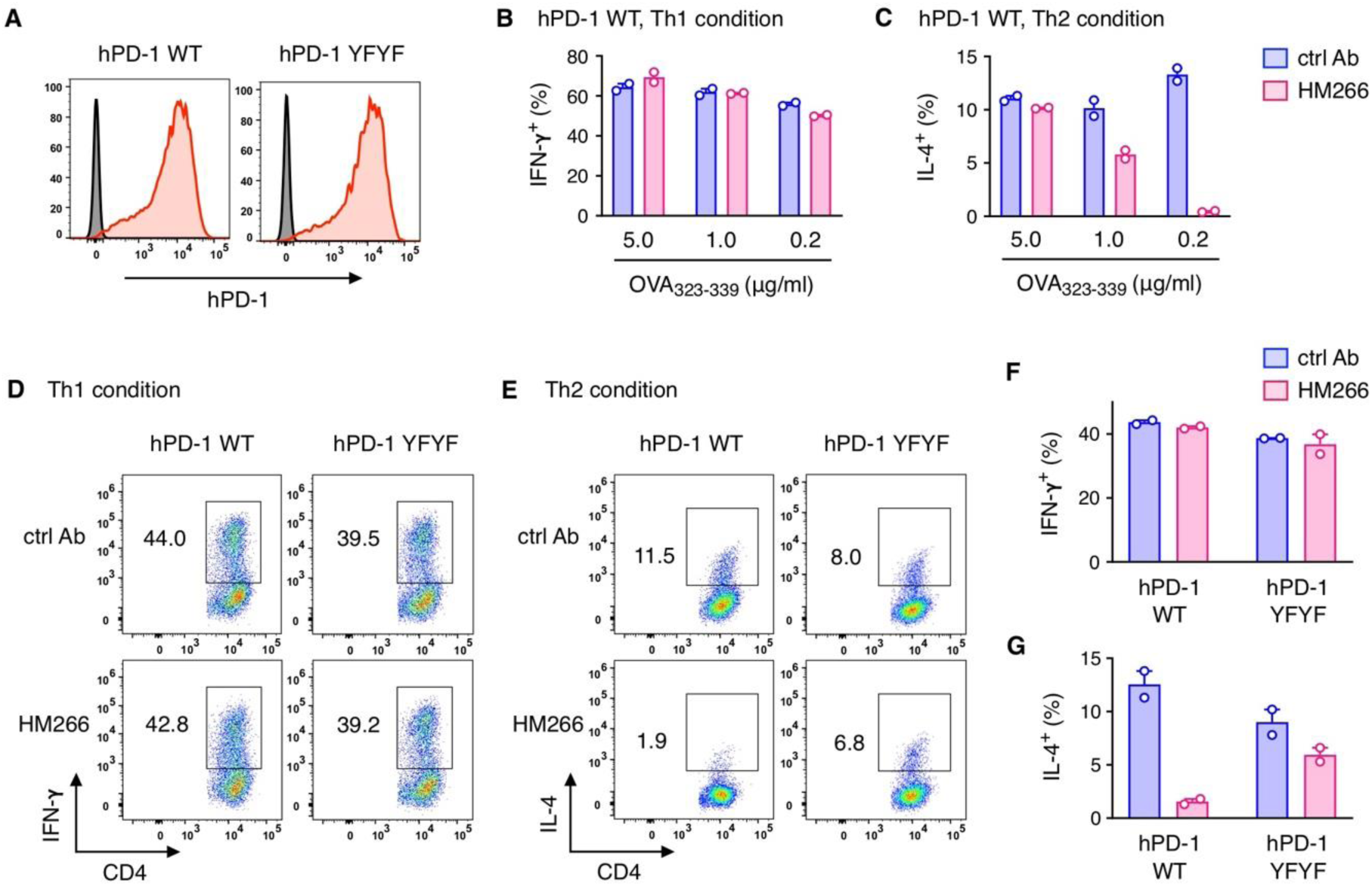
PD-1 agonist antibody strongly inhibits the induction of cytokine-producing Th2 cells. hPD-1-tranduced DO11.10 T cells were stimulated with OVA_323-339_ in Th1 or Th2- skewing condition using PD-L1-deficient A20 cells as APC. (A) hPD-1 expression in DO11.10 T cells after the transduction of wild-type hPD-1 or YFYF mutant. (B, C) The effects of anti-hPD-1 agonist mAb, HM266, on hPD-1 WT-transduced DO11.10 T cells in Th1 (B) or Th2-skewing condition (C). The concentrations of HM266 or isotype control were 5 μg/ml. Intracellular cytokine staining was performed on day 5. (D-G) The involvement of PD-1 signaling in Th2 inhibition by HM266. DO11.10 T cells expressing hPD-1 WT or hPD-1 YFYF were stimulated with OVA_323-339_ (1 μg/ml) in Th1 or Th2-skewing condition. Representative plots of intracellular cytokine staining on day 5 (D, E) and proportions of IFN-γ- (F) and IL-4-producing cells (G) within CD4^+^ cells. Data represent average ± SEM of duplicate samples.

The attenuated IL-4-producing Th2 cell development by early PD-1 stimulation was further confirmed by another anti-hPD-1 agonist antibody, J116, which has the less potent agonistic activity than HM266 (Figure 4A). J116 still impaired the induction of IL-4-producing cells, but not IFN-γ-producers, as shown by intracellular staining (Figure 4B-E) and by cytokine production in restimulated CD4^+^ T cells (Figure 4F, G). Again, the defect of PD-1 signaling in hPD-1 YFYF-expressing cells abolished the inhibitory effect of J116 on Th2 development.

**Figure 4.**
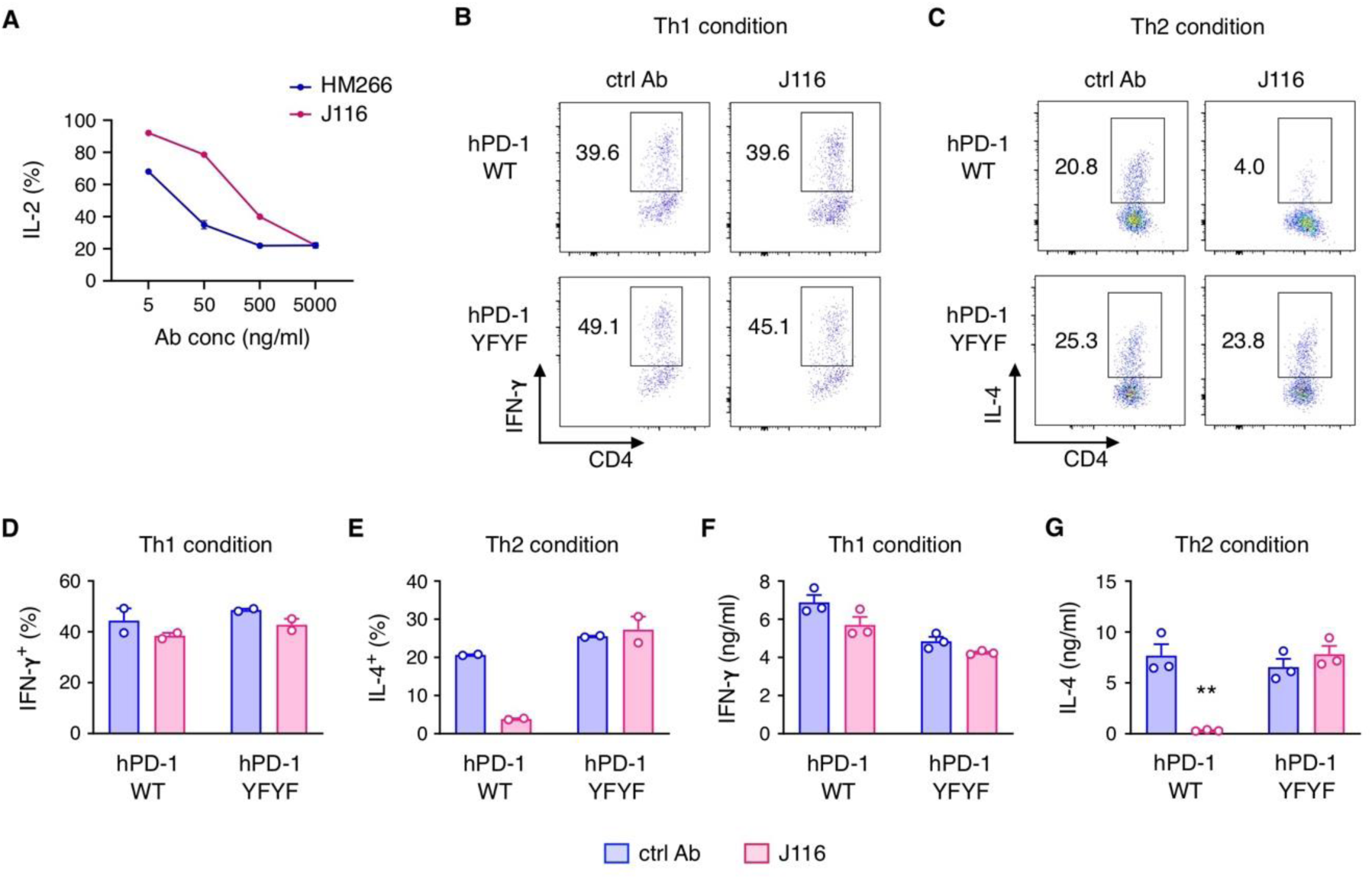
J116 inhibits the induction of IL-4-producing Th2 cells. (A) The immunosuppressive activity of anti-hPD-1 mAb J116. The PD-1 agonistic activity was determined by the inhibition of IL-2 production from DO11.10 T cell hybridoma. OVA_323-339_ peptide (2 μg/ml) was added to stimulate hPD-1-transduced DO11.10 cells in the presence of IIA1.6 cells that express murine FcγRIIB but lack PD-L1. (B-G) DO11.10 T cells were transduced with wild-type hPD-1 or YFYF mutant and skewed into Th1 or Th2 cells in the presence of J116 or mouse IgG1 isotype control (5 μg/ml). The concentration of OVA_323-339_ was 1 μg/ml. Intracellular cytokine staining was performed on day 5. Representative plots of cytokines staining (B, C) and percentages of IFN-γ- (D) and IL-4-producing cells (E) in CD4^+^ T cells. The same number of T cells were restimulated with anti-CD3 mAb on day 5, and IFN-γ- (F) and IL-4 levels (G) after 24 h were determined in the culture supernatant. Data represent average ± SEM of triplicate (A, F, G) or duplicate (D, E) samples. **p < 0.01; Student’s t-test.

Next, we examined more physiologically-relevant condition, which allows development of both Th1 and Th2 cells, instead of the skewing conditions inducing exclusively Th1 or Th2 cells. Sensitization of bone marrow-derived dendritic cells (BMDC) with house dust mite (HDM) extract has been known to induce preferential Th2-type immune response (*28*). The culture supernatant of HDM-treated BMDC was added to hPD-1-transduced DO11.10 cells. In this setting, similar proportions of IFN-γ- and IL-4-producers were observed in activated DO11.10 cells. HM266 was inhibitory to the development of both types of effectors, but the degree of inhibition was greater for IL-4-producers than IFN-γ-producers (Figure 5).

**Figure 5.**
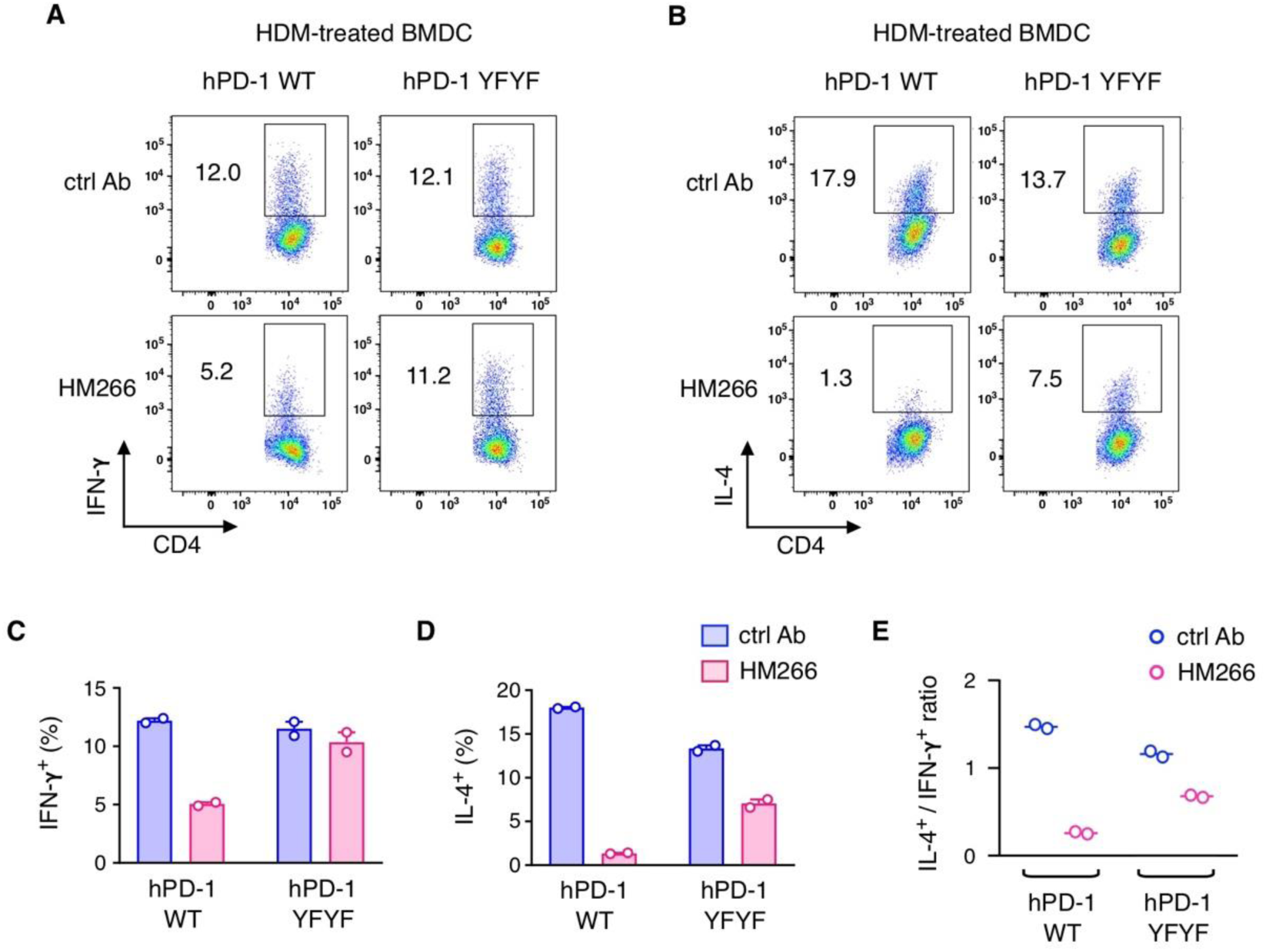
Immune deviation by PD-1 agonist antibody in a Th2-prone condition. hPD-1- tranduced DO11.10 T cells were stimulated with OVA_323-339_ (1 μg/ml) in the presence of HDM-stimulated BMDC supernatant using PD-L1-deficient A20 cells as APC. The concentrations of HM266 or isotype control were 5 μg/ml. (A, B) Representative plots of IFN-γ- (A) and IL-4-producing CD4^+^ cells (B) on day 5. DO11.10 T cells received transduction of either wild-type hPD-1 or YFYF mutant. (C, D) Proportions of IFN-γ- (C) and IL-4-producing cells (D) within CD4^+^ cells. (E) IL-4^+^ event / IFN-γ^+^ event ratios in individual samples. Data represent average ± SEM of duplicate samples.

The above experiments used hPD-1-transduced DO11.10 cells to test anti-hPD-1 agonist mAbs; therefore, those T cells express PD-1 at higher levels than that can be seen in spontaneous expression. To further confirm the attenuated Th2 development by PD-1 agonist at physiological levels of PD-1 expression, we produced hPD-1-knock-in (hPD-1-KI) DO11.10 mice, which express hPD-1 in place of mouse PD-1. These CD4^+^ T cells upregulated hPD-1 soon after the antigenic stimulation (Figure 6A, B). Stimulation of hPD-1 with HM266 in the early phase of functional differentiation diminished the development of Th2 cells, but not Th1 induction (Figure 6C, D). Cytokine levels from the restimulated T cells confirmed that T cell activation in the presence of PD-1 agonist mAb largely reduced IL-4/IL-13-producing Th2 cells, while IFN-γ-producing Th1 cells received only minor effect (Figure 6E-G). Taken together, PD-1 stimulation is inhibitory to the development of effector T cells, and its influence is substantial to Th2 immunity.

**Figure 6.**
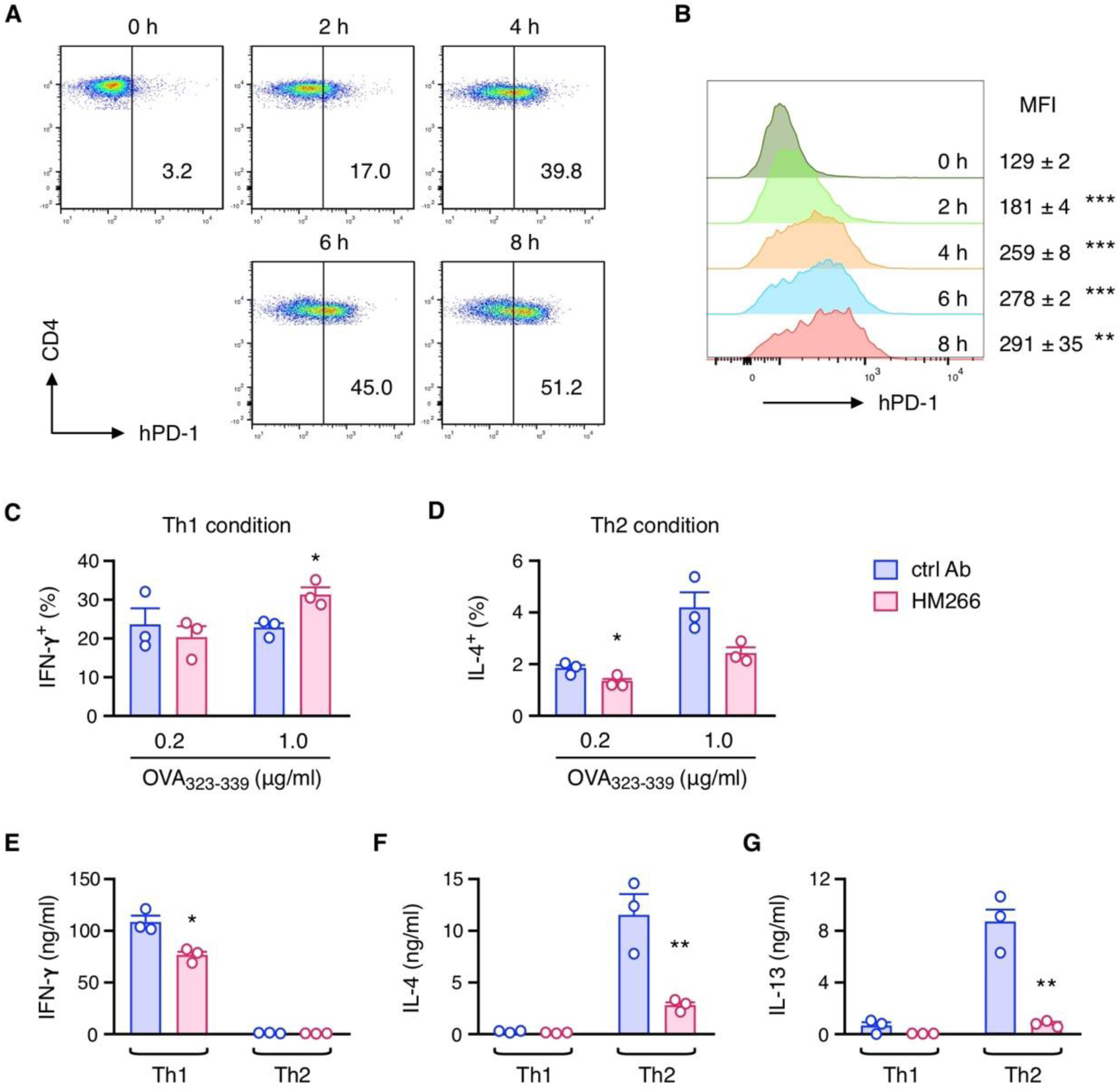
Spontaneous PD-1 upregulation in T cells sufficiently suppress functional differentiation into Th2 cells when stimulated with PD-1 agonist antibody. CD4^+^ CD62L^+^ cells from hPD-1-KI DO11.10 mice were stimulated with OVA_323-339_-pulsed PD-L1-deficient A20 cells. (A, B) Early PD-1 upregulation in the stimulated hPD-1-KI DO11.10 cells. OVA_323-339_ concentration was 1 μg/ml. Numbers indicate percentages of PD-1^+^ in CD4^+^ cells and mean fluorescence intensity of PD-1 (B). (C, D) The inhibition of Th2 cell induction by HM266, anti-hPD-1 agonist mAb. hPD-1-KI DO11.10 cells were stimulated with the indicated concentrations of OVA_323-339_ for 3 days. HM266 or control mouse IgG1 was added at 5 μg/ml. The activated CD4^+^ T cells were restimulated with immobilized anti-CD3 mAb for 6 h. IFN-γ-producing cells in Th1-skewing condition (C) and IL-4-producing cells in Th2-skewing condition (D) were evaluated by subsequent intracellular cytokine staining. (E-G) Cytokine-producing activities of activated CD4^+^ T cells. IFN-γ (E), IL-4 (F) and IL-13 (G) levels in the culture supernatant were evaluated after restimulation of activated T cells (2 x 10^5^ cells) on day 3 with immobilized anti-CD3 mAb for 24 h. Data represent average ± SEM of triplicate samples. *p < 0.05, **p < 0.01, ***p < 0.001; Student’s t-test.

### PD-1 agonist antibody suppresses allergic inflammation in vivo

Remarkable suppression of Th2 cell induction suggests that PD-1 is a potential target for the treatment of allergic diseases. To induce allergic asthma, we sensitized hPD-1-KI C57BL/6 mice with HDM extract. HDM extract is a potent allergen causing asthma in human and mice by the action of Th2-type cytokines such as IL-4, IL-5 and IL-13. Daily intranasal HDM challenge to the sensitized hPD-1-KI mice induced allergic inflammation as shown by massive infiltration of inflammatory cells in the lung. Eosinophils accounted for a large part of cell infiltrates in the bronchoalveolar lavage (BAL). CD4^+^ T cells in the inflamed lung and BAL expressed PD-1 along with CD69 and TIGIT (Figure 7, Figure 7-supplement 1).

**Figure 7.**
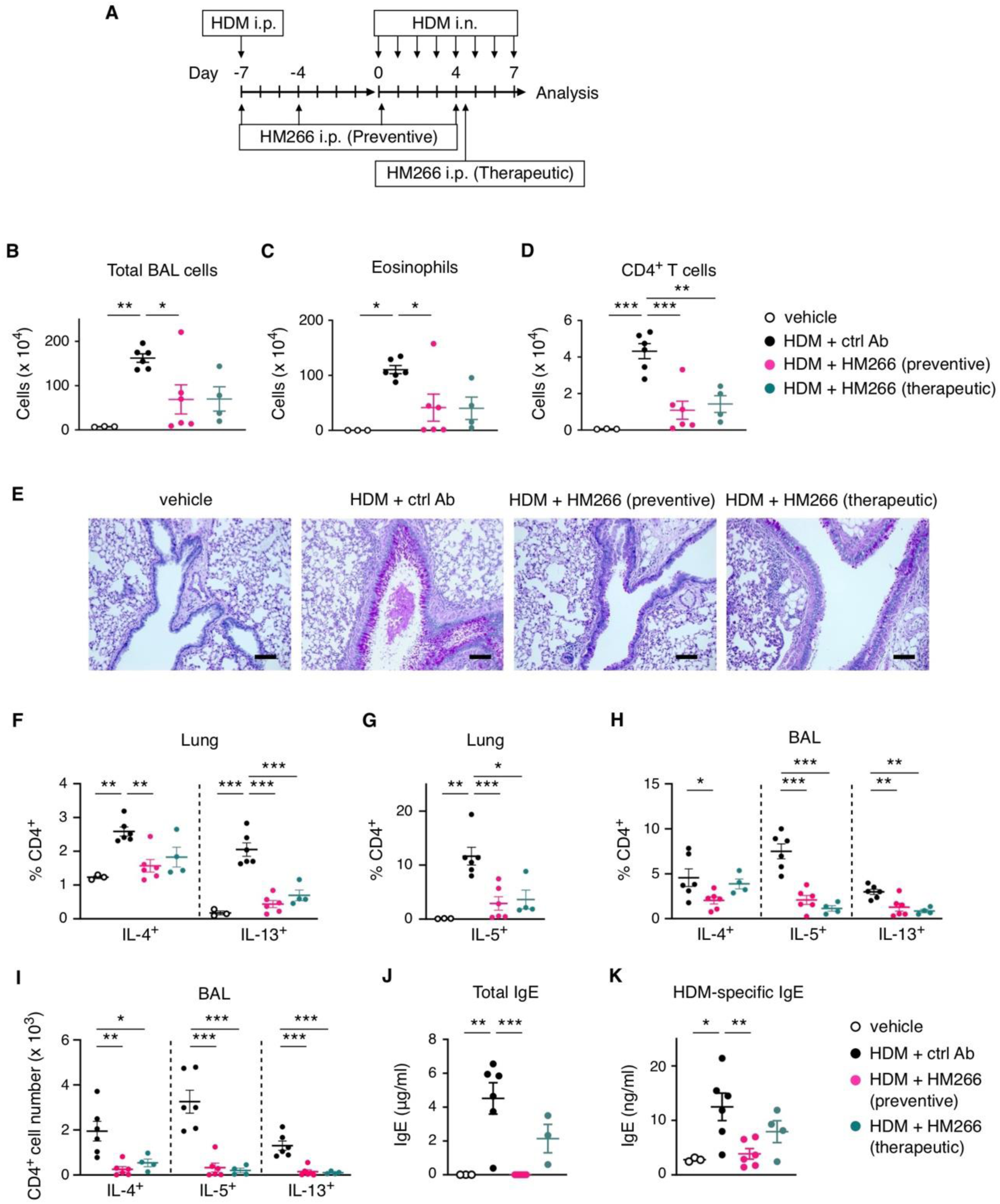
PD-1 agonist antibody ameliorates allergic inflammation. (A) The induction of allergic asthma in HDM-sensitized hPD-1 knock-in mice. Treatment with anti-hPD-1 agonist mAb, HM266, was done in two different schedules as indicated. (B-D) Infiltrated cell numbers in the BAL. Number of eosinophils (C) and CD4^+^ T cells (D) were calculated from flow cytometric analysis. (E) PAS staining of the lung sections. Scale bars represent 100 μm. (F-H) Intracellular cytokine staining of CD4^+^ T cells from the lung tissue (F, G) and BAL fluid (H, I). Panels indicate percentages in CD4^+^ T cells (F-H) and the numbers of cytokine-producing CD4^+^ T cells. (J, K) Plasma levels of total IgE (J) and HDM-specific IgE levels (K) on day 7. Data represent average ± SEM of vehicle control (n = 3), HDM + ctrl Ab (n = 6), HDM + preventive HM266 (n = 6) and HDM + therapeutic HM266 (n = 4) groups. *p < 0.05, **p < 0.01, ***p < 0.001; Tukey-Kramer test.

The efficacy of HM266 against allergic inflammation was examined in HDM-induced asthma model. When HM266 treatment started at the time of sensitization (Figure 7A, preventive setting), the PD-1 agonist strongly reduced infiltration of eosinophils and CD4^+^ T cells in the BAL (Figure 7B-D) and mucus hyperproduction in the lung (Figure 7E). Considerable portion of CD4^+^ T cells in the BAL and lung tissue were Th2-type cells producing IL-4, IL-5 and/or IL-13. HM266 significantly reduced proportions of these cytokine-producing cells (Figure 7F-H). With the drop in CD4^+^ T cell number taken into calculation, the treatment with PD-1 agonist strongly prevented the bronchoalveolar Th2 cell accumulation (Figure 7I). Serum IgE levels were significantly decreased by HM266 corresponding to the reduction in Th2 cytokines (Figure 7J, K). Of note, PD-1 stimulation was effective even when applied only after the elicitation of lung inflammation (Figure 7A, therapeutic setting). Single dose of HM266 after four consecutive intranasal HDM challenges could significantly reduce bronchoalveolar infiltration of eosinophils and CD4^+^ T cells, especially producers of IL-5 and IL-13 (Figure 7F-I). IgE levels after the therapeutic treatment with HM266 showed a decreasing trend, though it was not statistically significant. This result suggests that pharmacological PD-1 stimulation can alleviate allergic inflammation by blocking Th2-type immune response.

## Discussion

In this study, antigenic stimulation of T cells was found to quickly induce PD-1 expression on their surface. PD-1 stimulation during early stage of effector T cell development inhibited the production of functional effector cells; however, this inhibitory activity did not equally affect functionally differentiated CD4^+^ T cell subsets. Specifically, the co-inhibitory signal of PD-1 severely impaired Th2 cell development. Such an immune bias by PD-1 stimulation was not observed in studies using gene-deficient mice.

Mice lacking PD-1 or PD-L1 indicated highly active Th1-type immune response in Leishmaniasis (*29*) and Schistosomiasis (*30*). Associated with the enhanced Th1 immunity, PD-1-knockout mice have shown intense CD8^+^ T cell-dependent proinflammatory activities in anti-tumor response (*31–33*) and in contact hypersensitivity (*34, 35*). In contrast, these gene-deficient mice failed to exaggerate allergic inflammation in allergic asthma (*24*) and atopic dermatitis models (*21*). The predominance of Th1-type immunity in PD-1/PD-L1- deficienct mice might be stemmed from predisposed functional changes in immune cells. Indeed, treatment of wild-type mice with PD-1 blocking antibodies has provided different results from gene-deficient mice in the induction of allergic inflammation.

Experimentally, allergic diseases are often induced by sensitization of mice with an antigen and subsequent exposure of the target tissue with the same antigen. The blockade of PD-1 or PD-L1 during the challenge did not affect eosinophils activation or IgE increase (*20, 22*). However, PD-1/PD-L1 blockade during the sensitization phase significantly increased IL-4/IL-5/IL-13-producing cells in allergic conjunctivitis (*20*). In Schistosomiasis, PD-1 blockade was also shown to increase Th2 cells (*36*). Consistent with our study, these results using timely blockade approach suggested that PD-1 plays an important role in the regulation of Th2 cell induction.

Interestingly, the blockade of PD-L1 or PD-L2 has provided different results. While allergic inflammation did not change or rather attenuated in PD-L1-deficient mice (*21, 24*), PD-L2 deficiency exaggerated the induction of allergic asthma and dermatitis. Consistent with this difference, anti-PD-L2 blocking antibody induced stronger allergic inflammation than PD-L1 blockade (*20, 22*). The relative importance of PD-L2 in allergic inflammation may be due to its IL-4-inducible nature in contrast to IFN-γ-dependent induction of PD-L1 (*37, 38*). A complex regulation may be underlying with the PD-L2 involvement in allergic inflammation since PD-L2 was also suggested to affect Th1/Th2 balance in a PD-1-independent mechanism (*23, 39*).

To directly examine the role of PD-1 on Th functional differentiation, we compared DO11.10 T cell activation in the presence or absence of PD-1 stimulators. PD-1 stimulation is suppressive to the expansion of DO11.10 cells as previously reported (*40*), but CD4^+^ T cells that managed to proliferate in the presence of PD-1 stimulator were almost exclusively IFN-γ producers. PD-1 stimulation strongly impaired the development of IL-4-producing DO11.10 cells even in the Th2-skewing condition. Consistent with this view, cancer patients who have undergone PD-1 blockade for immunotherapy often experience eosinophilia (*16, 17, 41–43*), suggesting PD-1-dependent regulation of Th2 response in humans.

It remains to be elucidated why functional differentiation into Th2 is more sensitive to PD-1 stimulation than Th1 development. Extensive in vitro studies on Th differentiation have shown the importance of early IL-4 production in Th2 differentiation (*44*). The source of IL-4 has been investigated to identify the key player of Th2 induction. In T cells, the lack of IL-2 was shown to impair Th2 differentiation, and early IL-2 production was suggested to mediate IL-4 induction (*45, 46*). PD-1 stimulation with ligands or agonist antibodies downregulates IL-2 production from T cells (*25, 47*). However, IL-2 downregulation was unlikely to account for the PD-1-dependent inhibition of Th2 differentiation because IL-2 supplementation did not reverse the impaired development of IL-4-producing cells even in the Th2-skewing condition.

Since the relation of TCR signal strength to Th differentiation has been suggested, PD-1- dependent attenuation of TCR signal strength might have affected Th2 differentiation. Stimulation of naïve Th cells with low-intensity TCR signals preferentially induced Th2 cells in experiments using small Ag dose and low-affinity antigenic peptides (*46, 48–50*). However, in different settings, low antigen dose could result in preferential Th1 induction both in vitro and in vivo (*1, 51, 52*). In our study, the balance between immunostimulatory TCR signal and PD-1-dependent immunosuppressive signal significantly affected Th1/Th2 balance. PD-1 stimulation was previously reported to moderately modify the cytokine profile in favor of Th2 cells when T cells were stimulated with plate-bound anti-CD3 mAb and PD-L1-Fc (*53*). Although it has been frequently used to study PD-1-mediated immunosuppression, the co-immobilization of PD-L1-Fc sometimes causes PD-1-independent immunosuppression most likely due to the reduction of T cell engagement with anti-CD3 mAb (*25, 54*). The reason for the discrepancy with our result is unclear, but the difference in TCR signaling intensity might have influenced the outcome. Downstream TCR signaling, early ERK activation has been suggested to mediate Th2 differentiation (*55–57*). Inhibitory effect of PD-1 on ERK phosphorylation in the TCR signaling (*25*) may be involved in the mechanism of the attenuated Th2 differentiation. Interestingly, CTLA-4-Ig was previously shown to inhibit Th2 induction (*58*). The mechanism remains to be explored how co-stimulatory and co-inhibitory pathways influence the regulation of Th2 differentiation.

The overall role of PD-1 is suppressive to Th2 cells by inhibiting their establishment as well as effector functions, raising a possibility that PD-1 agonists are effective in the treatment of allergic diseases. We have previously identified anti-hPD-1 agonist antibodies that can stimulate the immunosuppressive activity of PD-1. The PD-1 agonist antibodies were effective in inhibiting various types of immune responses as shown in CD8^+^ T-dominant acute graft-versus-host disease and CD4^+^ T-dependent colitis induction (*25*). Other group also showed the alleviation of ILC2-induced lung inflammation (*59*) and neutrophilic lung inflammation by PD-1 agonist antibody (*60*). In our current study, the PD-1 agonist antibodies demonstrated anti-inflammatory efficacy to eosinophilic inflammation in allergic asthma. The treatment prevented IgE increase, eosinophilic infiltration and mucus production, which are correlated with the suppression of Th2 cytokines. Importantly, PD-1 agonist was effective even after the elicitation of disease. Taken together, PD-1 agonist antibody holds a promise as a therapeutic of various allergic disorders through the inhibition of type 2 immunity by targeting both Th2 cells and ILC2.

In conclusion, we found that the co-inhibitory signal of PD-1 not only reduces the intensity of effector T cell activities but also deviates functional differentiation of CD4^+^ T cells. The stimulation of PD-1 signaling strongly inhibited Th2 induction, and PD-L1 blockade resulted in a permissive condition that allowed Th2 abundance. The significance of PD-1-dependent regulation in Th2 response has been also suggested in human patients who received anti-PD-1 blocking Ab for cancer immunotherapy. Eosinophilia in these patients may indicate the sign of immunoenhancement by PD-1 blockade just as concurrent immune-related adverse events (irAE) has been found to correlate with better prognosis (*16, 17, 43*). Intense eosinophilic tissue inflammation sometimes complicates the anti-cancer treatment when PD-1 blockade induces severe allergic irAE such as drug reaction with eosinophilia and systemic symptoms (DRESS) syndrome and acute eosinophilic pneumonia (*18, 19*). PD-1 agonist Ab could regulate type 2 immunity to the opposite direction and prevent IgE increase and infiltration of eosinophils. The current study suggests the potential of PD-1-dependent immunoregulation as a target of anti-allergic treatment.

## Materials and Methods

### Mice

C57BL/6N mice were purchased from Japan SLC (Hamamatsu, Japan). DO11.10 TCR transgenic mice (Stock No: 003303) were purchased from the Jackson laboratory (Bar Harbor, ME). C57BL/6-backgroung hPD-1-knock-in mice (Accession No. CDB0117E: https://large.riken.jp/distribution/mutant-list.html) were generated as described previously (*25*). hPD-1 knock-in DO11.10 mice were produced by breeding DO11.10 mice with BALB/c-background hPD-1-knock-in mice. BALB/c hPD-1-knock-in mice (Accession No. CDB0119E: https://large.riken.jp/distribution/mutant-list.html) were generated by the insertion of hPD-1 cDNA-bovine poly A signal into the start codon of mouse PD-1 genome in BALB/c embryos as described for C57BL/6-backgroung hPD-1-knock-in mice (*25*). For the homologous recombination-mediated knock-in, the donor vector consisting of homology arms and hPD-1-bpA (bovine growth hormone poly A signal sequence) was generated to insert the hPD-1-bpA cassette at the 4 base upstream of the PAM sequence. gRNA site: 5’-GCC AGG GGC TCT GGG CAT GT-3’. Primer pairs for genotyping: wild-type FW 5’-AGG AGA CTG CTA CTG AAG GC -3’ and REV 5’-CCA ATC CGT GTA ACC AGG -3’ (245 bp); knock-in FW 5’-CAG GCC TCG ACA CCC ACC -3’ and REV 5’-CAG CCC AGT TGT AGC ACC -3’ (536 bp). All mice were maintained under specific pathogen free condition in the animal facility at Oriental BioService or RIKEN Kobe Branch. Both sexes were used for experiments. All animal experiment protocols were approved by the IACUC of Foundation for Biomedical Research Institute at Kobe (#20-04) and of RIKEN Kobe Branch (#A2017-08).

### Cell lines

A20 cells (RRID: CVCL_1940) were provided by Dr. Tasuku Honjo (Kyoto University). PD-L1-deficient A20 cells were produced by the following method. Guide RNA (target sequence: TCCAAAGGACTTGTACGTGG) was prepared using the GeneArt Precision gRNA Synthesis Kit (Invitrogen, catalog# A29377) and introduced into A20 cells together with GeneArt Platinum Cas9 Nuclease (Invitrogen, catalog# B25640) using the Neon Transfection System (Invitrogen). PD-L1-negative cells were isolated by several rounds of FACS sorting, followed by limiting dilution. Clone with the similar capability to stimulate T cells as parental A20 cell was chosen and used for further studies. PD-1 knockout DO11.10 T cell hybridoma and PD-L1 knockout IIA1.6 cells were obtained from Dr. Taku Okazaki (Tokyo University). Cell culture was done using RPMI 1640 (Gibco, catalog# 11875-093) supplemented with 10% fetal bovine serum (FBS), 6.25 mM HEPES, 2.5 mM L-glutamate, 0.625 mM sodium pyruvate, 0.625× nonessential amino acid solution, 62.5 μM 2-mercaptoethanol, penicillin (125 U/ml), streptomycin (125 μg/ml), and gentamycin (6.25 μg/ml) unless stated otherwise.

### Antibodies

Antibodies used in this study are listed in Table 1. Anti-hPD-1 mAbs, HM266 and J116, and anti-mouse PD-L1 mAb, 1-111A, were prepared from the culture supernatants of hybridoma cells. The hybridoma cells were grown in CD hybridoma medium (Gibco, catalog# 11279-023) supplemented with 8 mM L-glutamine, penicillin (20 U/ml) and streptomycin (20 μg/ml) using a CELLine bioreactor flask (Duran Wheaton Kimble, catalog# WCL1000). Antibodies in the culture supernatant were purified with Protein A-R28 (Ab-Capcher ExTra; Protenova, catalog# P-003).

**Table 1.**
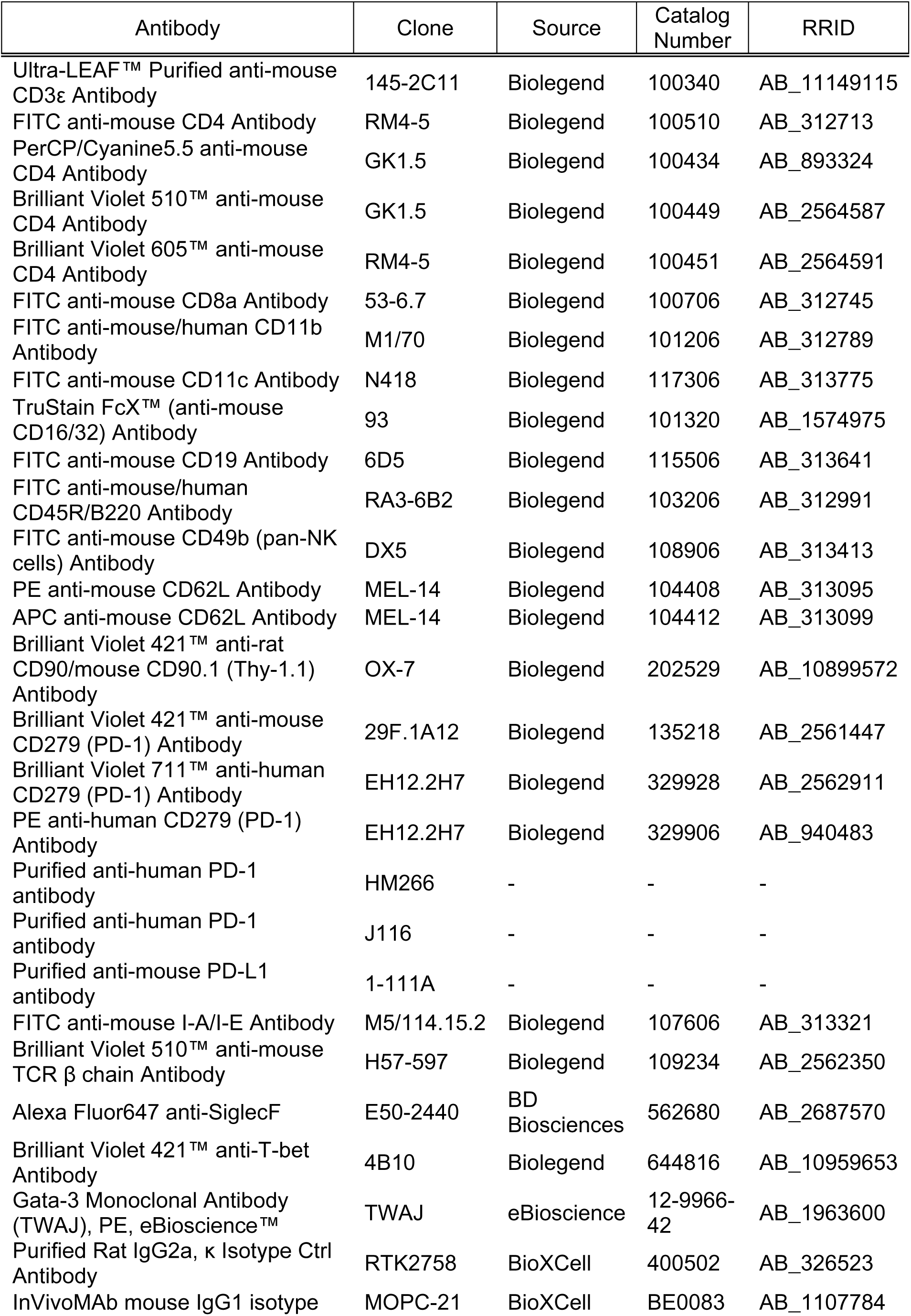
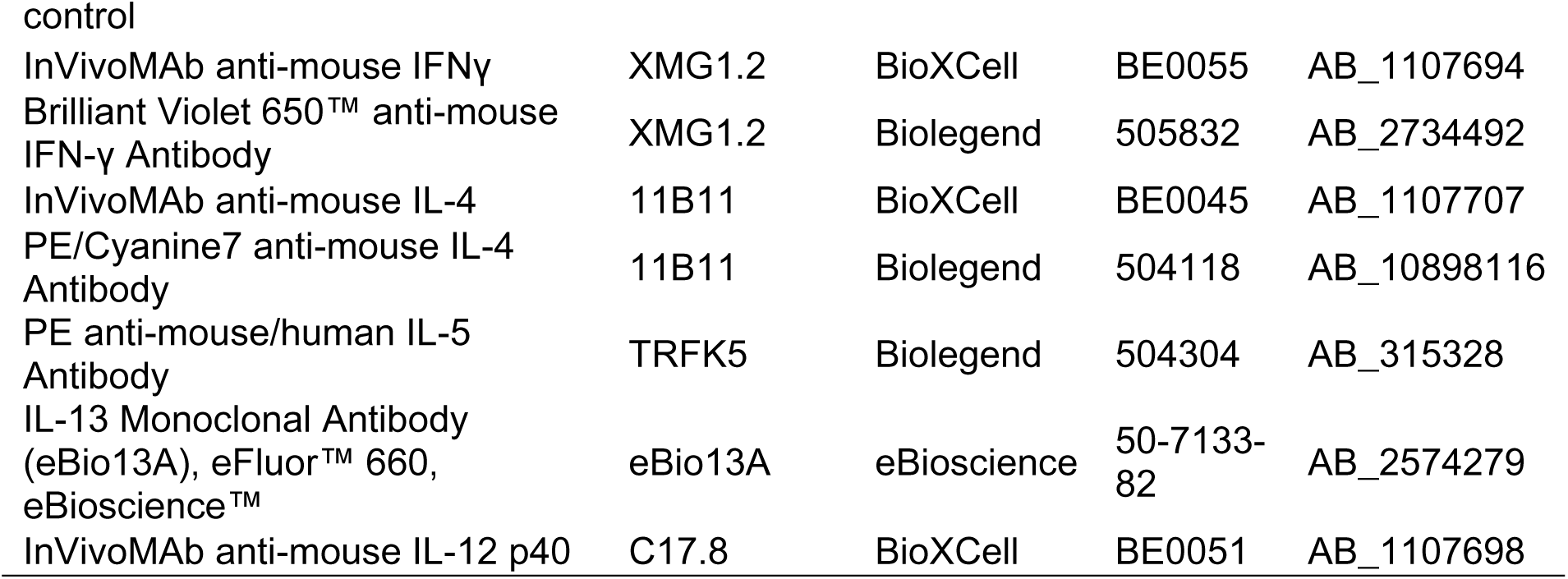
List of antibodies used in this study.

### Purification of naïve CD4^+^ cells

Naïve CD4^+^ T cells were sorted from DO11.10 splenocytes. After the lysis of red blood cells using ACK lysing buffer (Gibco, catalog# A10492-01), spleen cells were stained using FITC-labeled antibodies to CD8, CD19, CD45R/B220, CD11c, CD11b, CD49b and I-A/I-E. Antibody-labeled cells were depleted by labeling with anti-FITC MACS beads (Miltenyi Biotec, catalog# 130-048-701) and subsequent negative sorting using autoMACS Pro Separator (Miltenyi Biotec). The enriched CD4^+^ T cell fraction was further stained with PerCP-Cy5.5-labeled anti-CD4 mAb and APC-labeled anti-CD62L mAb, and CD4^+^ CD62L^+^ cells were purified using FACSAria or FACSMelody cell sorter (BD Biosciences).

### PD-1 expression

Purified DO11.10 CD4^+^ CD62L^+^ cells were stimulated with OVA_323-339_ peptide (Eurofins) using the same number of parental or PD-L1-deficient A20 cells as APC. The cell mixture was added in a 96-well flat-bottom plate, briefly spun down, and cultured in a CO_2_ incubator for the indicated period of time. PD-1 expression was analyzed by flow cytometry using BV421-anti-mouse PD-1 mAb or BV711-anti-hPD-1 mAb. Flow cytometry was performed by FACSFortessa X-20 (BD Biosciences) and analyzed with FlowJo software (BD Biosciences).

### T cell proliferation

Purified DO11.10 CD4^+^ CD62L^+^ T cells were labeled with CFSE (Invitrogen) in PBS at 5 μM for 30 min at 37 °C, followed by three rounds of washing by culture media prior to coculture with APCs. CFSE-labeled T cells were stimulated with OVA_323-339_ peptide in the presence of A20 cells. Time-dependent changes of CFSE signals of CD4^+^ cells were analyzed by flow cytometry.

### T helper subset differentiation

CD4^+^ CD62L^+^ T cells from DO11.10 mice were stimulated with OVA_323-339_ peptide using the same number of parental or PD-L1-deficient A20 cells. Prior to the co-culture, A20 cells were pretreated with 0.5 mg/mL mitomycin C (Fujifilm Wako Chemicals, catalog# 139-18711) at 37 °C for 2 hours and used after extensive wash. All functional differentiation experiments were conducted in the presence of IL-2 (2 ng/ml; Peprotech, catalog# 212-12). Additional cytokines or anti-cytokine antibodies were used for the specific skewing into Th1 or Th2 cells: IL-12p70 (1 ng/mL; Peprotech, catalog# 210-12), IFN-γ (1 ng/mL; Peprotech, catalog# 315-05) and anti-IL-4 mAb (5 μg/mL) for Th1; IL-4 (1 ng/mL; Peprotech, catalog# 214-14), anti-IFN-γ mAb (5 μg/mL), anti-IL-12 p40 mAb (5 μg/mL) for Th2. After 3 days, activated CD4^+^ T cells were restimulated with OVA_323-339_ in the same conditions for 1 or 2 more days. For the PD-L1 blockade experiment, anti-mouse PD-L1 mAb (5 μg/mL) or its isotype control mAb (RTK2758) was included in the culture. The effect of anti-hPD-1 agonist mAb, HM266 and J116 (5 μg/mL), was compared with that of isotype control mAb (MOPC21).

### Cytokine production

To evaluate cytokine-producing activity, DO11.10 cells after functional differentiation were washed with the cell culture media. The same number of activated CD4^+^ T cells (2 x 10^5^ cells) were restimulated with plate-bound anti-CD3 mAb on 96-well flat-bottom plates for 24 h. Cytokine levels in the culture supernatant were quantified by the use of Mouse IFN-γ DuoSet ELISA (R&D Systems, catalog# DY485), Mouse IL-4 DuoSet ELISA (R&D Systems, catalog# DY404) and Mouse IL-13 DuoSet ELISA (R&D Systems, catalog# DY413).

### Intracellular staining

Cells were stimulated with plate-bound anti-CD3 mAb on 96-well flat-bottom plates for 6 h in the presence of BD GolgiPlug (BD Biosciences, catalog# 555029) for the final 2 hours of culture. The stimulated cells were treated with anti-mouse CD16/32 mAb, followed by Fixable viability dye eFluor780 (eBioscience, catalog# 65-0865-14) and BV510-anti-CD4 mAb. Cell surface staining may include BV421-anti-Thy1.1 mAb when necessary. After fixation and permeabilization, cells were stained with BV650-anti-IFN-γ mAb, PE-anti-IL-5 mAb, PE-Cy7-anti-IL-4 mAb and eFluor 660-anti-IL-13 mAb. For the detection of transcription factors, cells stained with BV510-anti-TCRβ mAb and BV605-anti-CD4 mAb were fixed and permeabilized using True-Nuclear Transcription Factor Buffer Set (BioLegend, catalog# 424401) before staining with BV421-anti-T-bet mAb and PE-anti-GATA3 mAb.

### Retroviral transduction

hPD-1 retroviral plasmid was generated by inserting hPD-1 cDNA into MSCV-IRES-Thy1.1 DEST (Addgene, catalog# 17442). Point mutations were introduced by using QuikChange ll site-directed mutagenesis kit (Agilent, catalog# 200523). For bicistronic transduction of hPD1/GFP fusion and mSHP2-C459S protein, IRES-Thy1.1 sequence of MSCV-IRES-Thy1.1 was replaced with hPD-1/GFP-P2A-mSHP2-C459S sequence. The retroviral plasmids were transfected into Plat-E cells (Cell Biolabs, RRID: CVCL_B488) using Fugene HD Transfection Reagent (Promega, catalog# E2311). After 2-3 days, retroviral supernatant was pooled and passed through 0.45 μm Minisart Syringe Filter (Sartorius, catalog# 16533). The retrovirus supernatant was added to Retronectin (Takara Bio, catalog# T100A)-coated culture plate and centrifuged at 32 °C for 2 h. Purified naïve DO11.10 CD4^+^ T cells as described above were stimulated using Dynabeads Mouse T-Activator CD3/CD28 (Gibco, catalog# 11452D) for 24 h. Activated T cells were placed on retrovirus-coated culture plate and centrifuged at 800 x g for 10 min at 32 °C. This procedure was repeated on the next day, and hPD-1-transduced T cells was used on day 3 for further studies. Cellular expression of hPD-1 was confirmed by flow cytometry using PE-anti-hPD-1 mAb.

### Agonistic activity of anti-hPD-1 mAbs

hPD-1-expressing DO11.10 T cell hybridoma cells (5 × 10^4^ cells) were stimulated with OVA_323–339_ (2 μg/ml) presented on hPD-L1^−^ mFcγRIIB^+^ IIA1.6 B lymphoma cells (1 × 10^4^ cells). Culture supernatants were collected after 18 h, and IL-2 levels were determined using mouse IL-2 DuoSet ELISA (R&D Systems, catalog# DY402).

### Sensitization of BMDCs with HDM extract

Bone marrow cells from the femurs were treated with ACK lysis buffer and resuspended in the cell culture media. After incubating in a culture plate for 4 h at 37 °C, loosely adherent cells were collected and further cultured in media supplemented with recombinant mouse GM-CSF (20 ng/mL; Peprotech, catalog# 315-03). The same fresh GM-CSF-supplemented media was supplied on day 2 and 4. On day 6, the cells were further incubated in media without GM-CSF for 1 day before using them as BMDCs. HDM extract (Dermatophagoides pteronyssinus; Greer Laboratories, catalog# B70) was given to cultured BMDCs at 1 μg (total protein)/ml, and the supernatant was collected after 24 h.

### HDM extract-induced allergy model

hPD-1-KI C57BL/6 mice were randomly assigned to groups and were sensitized with HDM extract (10 ng Der p 1/mouse i.p.). A week later, intranasal challenge with HDM extract (25 μg of whole protein/25 μL per mouse) started and repeated for 7 consecutive days. The mice were euthanized 4 h after the last challenge. The BAL fluid was collected by introducing 1 ml PBS through the trachea. The lungs were processed with Lung Dissociation Kit (Miltenyi Biotec, catalog# 130-095-927) and GentleMACS Dissociator (Miltenyi Biotec) according to the manufacturer’s instructions, and the lung mononuclear cell were isolated by density centrifugation using 40 and 70% Percoll (Cytiva, catalog# 17089102). For the detection of eosinophils, BAL cells were incubated with anti-mouse CD16/32 mAb to block Fc receptors, and stained with FITC-anti-CD11c mAb, Alexa Fluor 647-anti-SiglecF mAb and Fixable viability dye eFluor780. Eosinophils were defined as CD11c^-^ SiglecF^+^ population by flow cytometry. For intracellular cytokine staining, BAL cells were stimulated with plate-bound anti-CD3 mAb on 96-well flat-bottom plates for 6 h in the presence of BD GolgiPlug for the final 2 hours of culture and were stained as described above.

### Histology

Lungs from HDM-induced allergic mice were collected and fixed in 10% neutral-buffered formalin. Fixed lungs were then processed for PAS staining by Applied Medical Research Laboratory.

### Plasma IgE levels

Levels of total IgE and HDM-specific IgE in the plasma were determined by using ELISA MAX™ Deluxe Set Mouse IgE (Biolegend, catalog# 432404) and Mouse Serum Anti-HDM Dermatophagoides pteronyssinus IgE Antibody Assay Kit (Chondrex, catalog# 3037) according to manufacturer’s instructions.

### Statistics

Data are expressed as mean ± SEM of biological replicates. Data shown in Figures are representative of 2-3 independent experiments with similar results. Statistical significance was calculated by two-tailed Student’s t-test for two-group comparison and Tukey-Kramer test for multiple comparison. P values less than 0.05 were considered significant.

## Data Availability

The data are available from the corresponding author upon reasonable request.

## Author Contributions

Conceptualization: MT, TH, AO

Methodology: MT, AO

Investigation: MT, NI, YN, KS, YT, PL, AO

Resources (hPD-1-knock-in mice): HK

Writing – Original Draft: MT, AO

Writing – Review and Editing: MT, NI, YN, TH, AO

Supervision: TH, AO

Funding Acquisition: TH

## Acknowledgements

This work was supported by Meiji Seika Pharma Co. Ltd. and the Foundation for Biomedical Research and Innovation at Kobe. The authors thank to Miwa Okada, Kumiko Yonezaki, Yukako Kamita and Chiyomi Ito for the antibody production and mouse colony management. Anti-hPD-1 agonist antibodies were previously developed in our collaboration with Drs. Satoshi Nagata and Haruhiko Kamada (National Institutes of Biomedical Innovation, Health and Nutrition).

---

**Figure 7, supplement 1.**
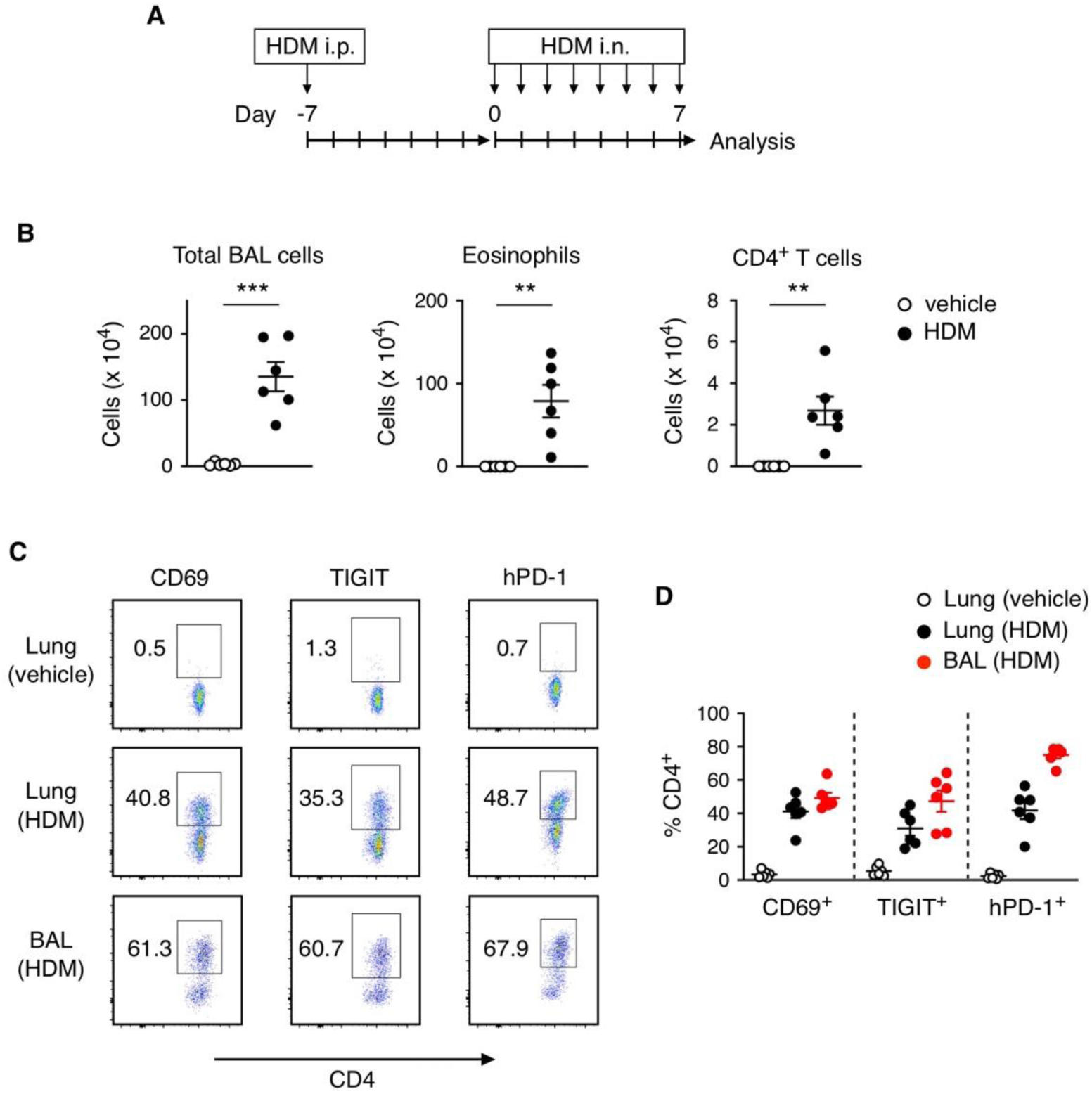
PD-1 expression on CD4^+^ T cells from allergic hPD-1 knock-in mice. (A) HDM extract induced lung allergy was induced on hPD-1 knock-in mice as indicated. Mice was euthanized 4 h after the final intranasal challenge. (B) BAL cell numbers. Numbers of eosinophils and CD4^+^ T cells were calculated from flow cytometry analysis. (C) Representative FACS plots of CD69, TIGIT, and hPD-1 expression on CD4^+^ T cells collected from BAL or lung cells. (D) Percentages of CD69^+^, TIGIT^+^, and hPD-1^+^ cells in CD4^+^ T cells. Data represent average ± SEM of 6 mice. **p < 0.01, ***p < 0.001; Student’s t-test.

